# Digitally Predicting Protein Localization and Manipulating Protein Activity in Fluorescence Images Using Four-dimensional Reslicing GAN

**DOI:** 10.1101/2022.07.24.501328

**Authors:** Yang Jiao, Mo Weng, Lingkun Gu, Yingtao Jiang, Mei Yang

## Abstract

**Motivation:** While multi-channel fluorescence microscopy is a vital imaging method in biological studies, the number of channels that can be imaged simultaneously is limited by technical and hardware limitations such as emission spectra cross-talk. One feasible solution is using deep neural networks to model the localization relationship between two proteins so that the localization of a protein can be digitally predicted. Furthermore, the input and predicted localization implicitly reflects the modeled relationship. Accordingly, observing the predictions via repeatedly manipulating input localizations is an explainable and feasible way to analyze the modeled relationships between the input and the predicted proteins.

**Results:** We propose a Protein Localization Prediction (PLP) method using a cGAN named Four-dimensional Reslicing Generative Adversarial Network (4DR-GAN) to digitally generate additional channels. 4DR-GAN models the joint probability distribution of imaged and target proteins by simultaneously incorporating the protein localization signals in four dimensions including space and time. Because protein localization often correlates with protein activation state, with accurate PLP, we further propose two novel tools: digital activation (DA) and digital inactivation (DI) to digitally activate and inactivate a protein and observe the response of the predicted protein localization. Compared with genetic approaches, these tools allow precise spatial and temporal control. A comprehensive experiment on four groups of proteins shows that 4DR-GAN achieves higher-quality PLP than Pix2Pix and the DA and DI responses are consistent with the known protein functions. The proposed PLP method helps simultaneously visualize additional proteins and DA and DI provide guidance to study localization-based protein functions.

**Availability and Implementation:** The open-source code is at https://github.com/YangJiaoUSA/4DR-GAN.

## 1. Introduction

Fluorescence microscopy, where samples are labeled with fluorescent probes (a.k.a. fluorophores), is one of the most versatile optical imaging methods. It allows visualization and quantitation of various aspects of the target proteins, including their level, localization, behavior, and interaction with other proteins. Usually, each protein of interest is labeled by one type of fluorophore, and its signals are collected as one channel. In laser-scanning confocal microscopy, a labeled protein can be imaged in a three-dimensional volume with three spatial axes. Application to live tissues or organisms generates time-lapse data sets with the time axis. Further, if multiple proteins are labeled and imaged, additional channels are obtained, which result in information-rich five-dimensional data sets.

Although multi-channel imaging is a powerful tool to understand protein functions, in practice, the number of proteins that can be imaged simultaneously is limited. This is because the emission spectra of individual fluorophores is often too wide to be sufficiently separated. Additionally, the choice of fluorophores is limited by the quantum yield and photostability of fluorophores^1^, as well as the in vivo concentration of target proteins. When it comes to live imaging, the choice of fluorophores is particularly limited because signals from live samples are much weaker. These limitations contribute to the difficulties of simultaneously imaging more than two proteins in live samples. Beyond the choice of compatible fluorophores, the number of channels is also bound by the availability of laser lines and detectors of a microscope, the demand for acquisition speed, and the availability of genetically labeled proteins. Without a proper tool, simultaneously observing and even studying multiple proteins has been quite a challenge.

One feasible way to alleviate this challenge is to use machine learning methods to digitally predict the localization of unimaged proteins, using the localization information obtained from the imaged proteins. As a promising candidate model for this task, conditional generative adversarial networks^2^ (cGANs) are able to take an input image and generate the desired output image. A cGAN usually has a generator and a discriminator that are both convolutional neural networks. The generator uses network parameters to implicitly model the joint probability distribution of the inputs and the outputs so that it generates the desired output for any given new input. Theoretically, if enough samples and training time are offered, the modeled probability distribution can match the true distribution^3^. The discriminator models the conditional probability distribution of real and fake sample judgement given the real and generated protein localization. When given a new input, the generator tries to produce an output that to fool the discriminator. In biological image processing, cGANs are popular in multiple topics including data augmentation^4–7^, domain translation^8,9^, resolution enhancement^10–13^, virtual stain^14–19^, stain normalization^20,21^, among others ^22–24^. Particularly, Pix2Pix^25^ is a successful example of cGANs that shows effectiveness on multiple tasks such as image colorization and style transfer. A recent work^26^ attempted to predict the localization of a protein using another protein from 2D fluorescence images with Pix2Pix. However, this work failed to obtain pixel-wise accurate results, likely because it only considered the 2D correlation between proteins. In reality, proteins exist in three-dimensional space and are often temporally correlated because of reversible interaction and changing activities.

To address this caveat, we propose a *Protein Localization Prediction* (PLP) method using a new cGAN named Four-dimensional Reslicing Generative Adversarial Network (4DR-GAN). 4DR-GAN models the joint probability distribution of imaged and unimaged proteins by incorporating the correlations between two protein localizations manifested in four dimensions, three in space and one in time. To our knowledge, this is the first work on applying cGANs to 4D information modeling. The generator of the 4DR-GAN is an end-to-end network that takes a 4D image as an input and incorporates the spatial and temporal information by two encoding paths. Subsequently, the 5D feature maps are extracted from the two paths, and they are resliced to the same shape to be paired in space and time. The paired features are reconstructed to produce a 4D image as the input. In a nutshell, this 4DR-GAN, enables accurate prediction of protein localization that cannot be imaged together.

Furthermore, with the new capability of accurate PLP of fluorescence images, it opens the door to digitally manipulate a protein’s localization and activation, which is likely to trigger another protein response in localization prediction and reveal the protein relationships. In this regard, we further propose two novel tools, digital activation (DA) and digital inactivation (DI). DA digitally increases protein localization or protein activity, while DI performs in contrast.

DA and DI present advantages when compared with genetic knockout and knockdown in terms of protein activity manipulation. Essential in testing the function of a gene, genetic knockout removes the gene from the genome, and gene knockdown stops or decreases the expression of the targeted genes. However, gene knockout and knockdown have drawbacks that mostly originate from their limited spatial and temporal control capabilities. Applying genetic knockdown or knockout to undesired tissues or stages often complicates the analysis of gene functions. Another drawback is that these genetic approaches are unable to manipulate gene functions at subcellular levels, which is an important capability in understanding the differential protein functions at multiple subcellular compartments. In contrast, DA and DI can manipulate gene functions with precise spatial and temporal control and induce immediate effects, allowing gene function to be digitally removed or activated in any cells and subcellular regions at any time point. If the protein manipulation consistently leads to changes in prediction, the changes reflect the local or global relationship between the input and the predicted proteins, making DA and DI desirable tools for protein functional relationship study.

To test the effectiveness of PLP along with DA and DI, we used five-dimensional data sets from live imaging of Drosophila embryos that revealed the localization of two proteins in separate channels. These data sets offer rich temporal information since the subcellular localization of proteins changes rapidly in a developing embryo. The high spatial and temporal resolutions of these datasets (pixel size: ∼0.1 um; frame rate: 10 sec) allow us to test prediction accuracy at subcellular levels. The proteins involved are well studied in their localizations and functions, and therefore, they offer a variety of evaluation criteria.

We summarize our contributions in the following three aspects:

1. To visualize more proteins simultaneously in fluorescence microscopy, we propose a protein localization prediction (PLP) method to predict the localization of unimaged protein from imaged proteins using Four-dimensional Reslicing Generative Adversarial Networks, a new cGAN developed solely for this work. 4DR-GAN can simultaneously incorporate four-dimensional information for the purpose of protein localization prediction.
2. Based on PLP, we develop two new tools to digitally manipulate protein localization and activation: digital activation and digital inactivation. These tools allow precise spatial and temporal manipulation and induce an immediate response. A consistent response could provide clues to the functional relationship between the two proteins.
3. A comprehensive experiment on four groups of PLP shows the effectiveness of 4DR-GAN and the success of PLP. Compared with the existing network, the protein localization and dynamic behavior in our prediction results are closer to the ground truth. Through performing DA and DI on multiple groups of proteins, we obtained responses in the predicted protein localizations that are consistent with the known protein functions.

## 2. Materials and methods

### 2.1 Four-dimensional Reslicing GAN

The fundamental role of 4DR-GAN is to incorporate spatial and temporal information simultaneously in the input 4D image and produce realistic 4D output. In the case of PLP, by taking the channel of one protein localization as input, 4DR-GAN predicts that of another protein localization as output. 4DR-GAN consists of a 4D-reslicing generator (*G*) that predicts the protein localization in 4D images, and a 4D-consistency discriminator (*D*) that assesses the realness of the prediction in terms of localization, temporal consistency, and the input-target correlation. Fig. 1 demonstrates the structure of 4DR-GAN, where an input 4D image is denoted as 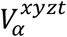, and a target 4D image denoted as 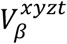. 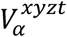 is first resliced into XYZ-T view as 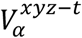 which sees the 4D image as XYZ-volumes with *t* frames, and XYT-Z view as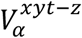, which sees it as XYT-volumes with *z* frames.

**Fig. 1.**
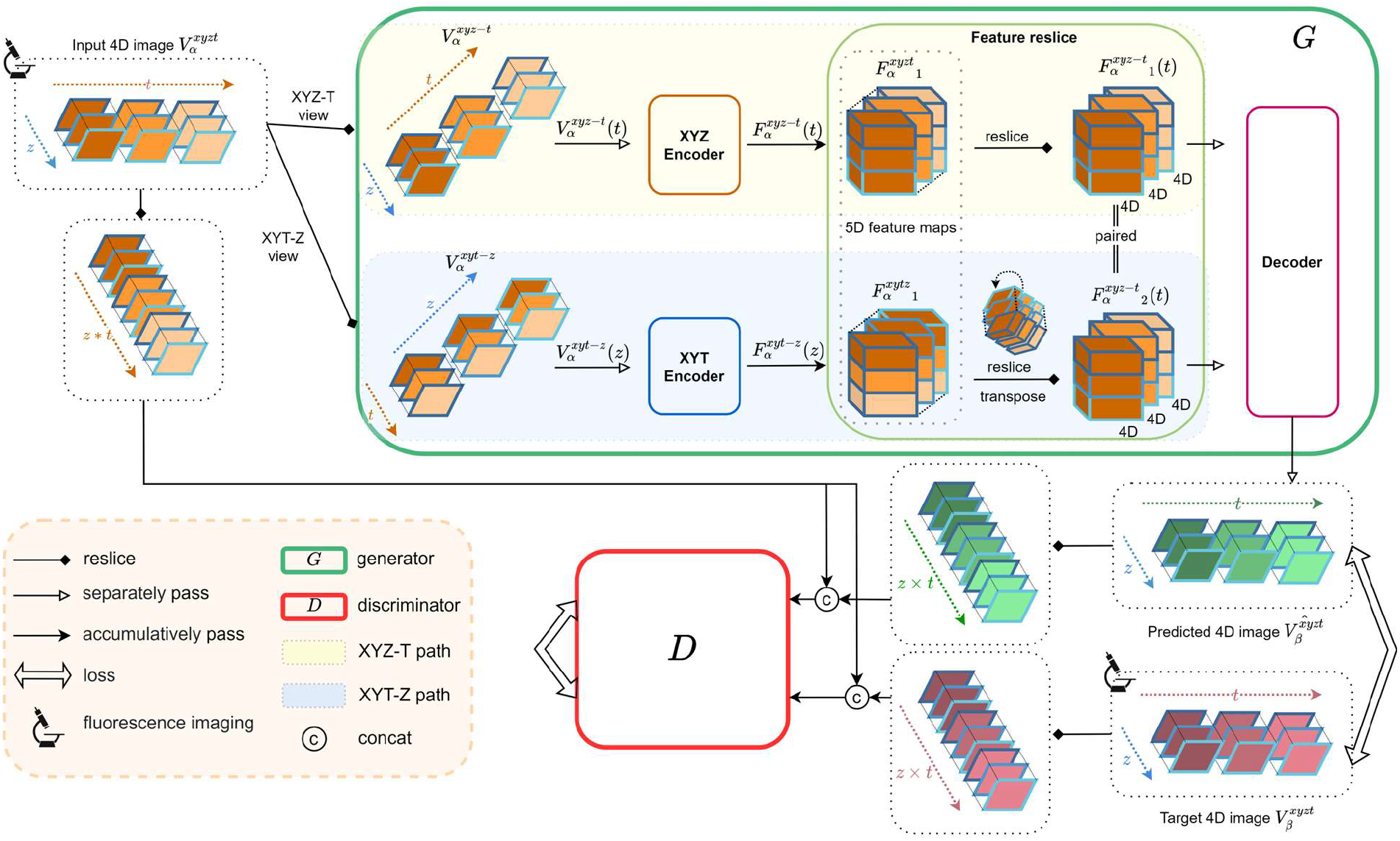
4D fluorescence image prediction by Four-dimensional Reslicing GAN (4DR-GAN). The overall flow of training the generator and the discriminator of 4DR-GAN. The input and the target 4D images constitute two channels of a 5D fluorescence image, which visualize the localizations of two proteins in the X, Y, Z, and T-axes. *G* is a dual-path network that separately encodes the XYZ-axis and XYT-axis information of the input 4D image. *D* justifies the realness of the predicted image by taking the 4D images that are resliced into XY(ZxT) view. Various types of arrows are used to distinguish different operations, as shown in the legend. Network implementation detail can be found in Supplementary.

Correspondingly, the generator *G* has two paths. In the XYZ-T path of *G*, the XYZ-volume of each *t* frame 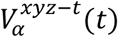 is sent to XYZ Encoder to obtain the feature maps, denoted as 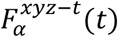, while in the XYT-Z path, the XYT-volume of each *z* frame is sent to XYT Encoder to obtain the feature maps, denoted as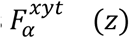. 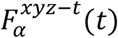 and 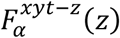 are 4D maps that incorporate both spatial and temporal information. All the feature maps, 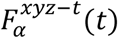 and 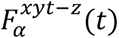, are further assembled into 5D feature maps denoted as 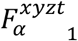 and 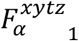, respectively. Subsequently, taking into account that 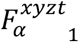 and 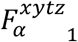 represent different views of the image, we reslice 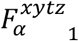 according to XYZ-T view to become 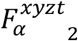, *which* spatially and temporally matches 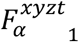. This reslicing operation is detailed in Method Section 2.

To reconstruct a 4D output, the two 5D feature maps 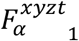 and 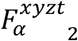 in *XYZ-T* view will be independently decoded to obtain 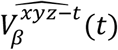 in all *t* frames. Specifically, the 5D feature maps 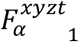 and 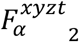 are resliced into individual 4D feature maps 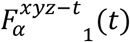 and 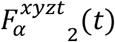 in *t* frames, and the corresponding pairs in time are sent to the Decoder to reconstruct the XYZ-volume 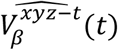. Upon the completion of reconstruction, the prediction in all *t* frames 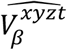 *is obtained*.

The discriminator *D* justifies the realness of the prediction by taking the 4D images that are resliced into XY(ZxT) view such as 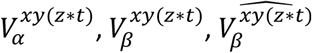. *D* takes the localizations of the input and the target proteins simultaneously, such as 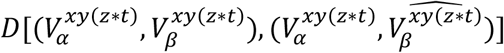. In this way, *D* justifies the localization and the temporal consistency of proteins, as well as the interaction between proteins.

### 2.2 Data acquisition

We applied this 4DR-GAN to the 5D data sets collected from live imaging of Drosophila early embryos involving three proteins: Myosin (Myo), E-Cadherin (E-Cad) and Ajuba (Jub). Embryos were dechorionated in 4% sodium hypochlorite, washed in water, and mounted in glass-bottom Petri dishes by the natural affinity between the vitelline membrane and the glass. The dish chamber was then filled with water and covered by an oxygen-permeable membrane. The imaging was performed with a Zeiss LSM 800 confocal microscope equipped with high sensitivity GaAsp detectors. The 488-nm and 561-nm lasers were used to excite GFP and mCherry, respectively. Images were acquired using a plan-Apochromat 63x/1.40 oil objective with the pinhole set at 1 Airy unit and the pixel size set at 0.124 µm. The z-stacks start from the embryo surface to 7 μm deep with 0.5-μm increments. The time interval between stacks is 10s.

The original 5D fluorescence images have two channels and slight variations in shapes. The two channels are split as input and target, and then resized and cropped into training and testing samples. Accordingly, we performed four groups of PLP: from Myo to E-Cad, from E-Cad to Myo, from Jub to Myo, and from Jub to E-Cad.

For each group of proteins, 516 samples with a size of 256×256×16×10×2 were used for training. Each sample had 256 pixels on x-axis and y-axis, 16 pixels on z-axis, and 10 time frames. Meanwhile, two channels were involved in each sample, where one channel was used as input, and another channel was as the target output (GT). These samples were cropped from twelve 5D fluorescence images. For validation, 129 samples were cropped from another three images. Three images containing the Myo localization with a size of 256×256×16×40×1 were used for testing.

## 3. Results

### 3.1 4DR-GAN generates high-quality localizations that have better FID scores than the compared baseline

To quantitatively evaluate the quality of the predictions generated by 4DR-GAN, we first employed the Fréchet Inception Distance (FID)^27^. It is a widely used metric that reflects the human perception of similarity because it employs a deep CNN layer closer to output nodes that correspond to real-world objects. In contrast, the traditional pixel-level metrics, such as Mean Square Error (MSE), Structural Similarity Index (SSIM), and Peak Signal-to-noise Ratio (PSNR), are mismatched with human perceptual preference^28^. Two widely used pretrained CNNs are employed in our evaluation: InceptionV3^29^ trained on ImageNet for image classification, and I3D^30,31^ trained on Kinetics400 for video recognition. Because our prediction output is 4D, it is sliced on the Z-axis and T-axis into 2D images to fit InceptionV3 so that the similarity between prediction output and ground truth was evaluated in XY-plane. To fit I3D, the prediction output is sliced along Z-axis or T-axis, which results in XYZ-volume and XYT-volume respectively. This allows the evaluation of volumetric and temporal consistency. We compared 4DR-GAN against Pix2Pix which was used in a previous work^26^, and we further developed Pix2Pix from 2D to 3D to optimize its ability in PLP.

Table 1 demonstrates the FID evaluation on the prediction of four groups of samples. For the prediction quality in 2D, 4DR-GAN surpasses Pix2pix as reflected on the FID with InceptionV3.

**Table 1.**
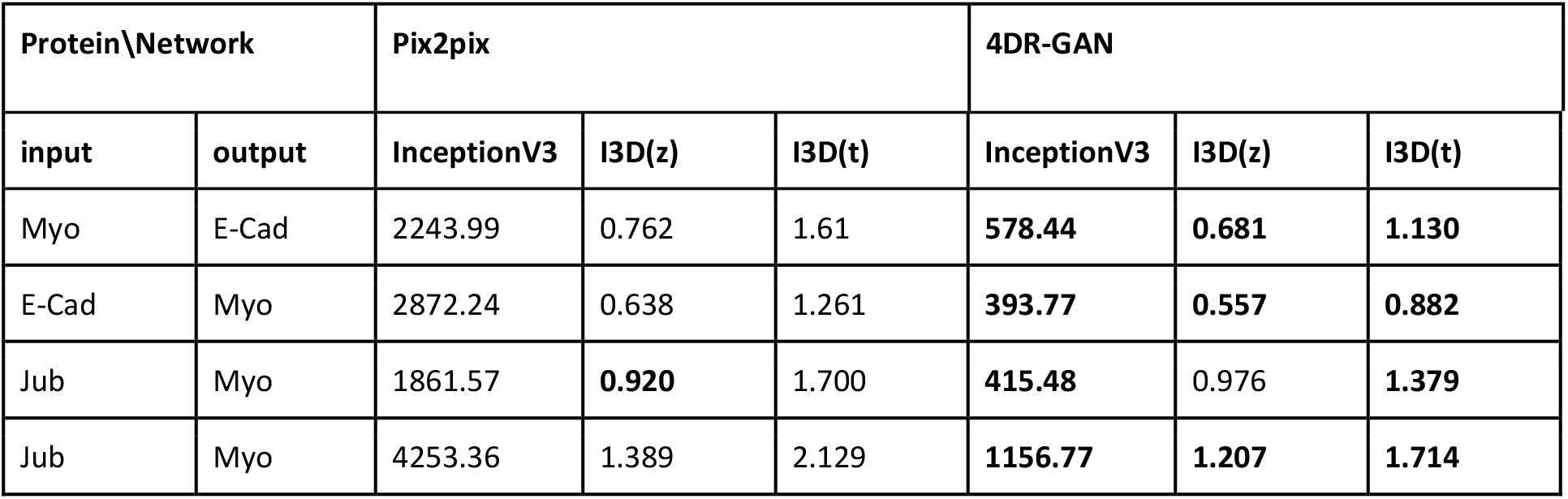
Prediction quality evaluation by FID score. Lower is better. The best results are in bold.

| Protein\Network |  | Pix2pix |  |  | 4DR-GAN |  |  |
| --- | --- | --- | --- | --- | --- | --- | --- |
| input | output | InceptionV3 | I3D(z) | I3D(t) | InceptionV3 | I3D(z) | I3D(t) |
| Myo | E-Cad | 2243.99 | 0.762 | 1.61 | **578.44** | **0.681** | **1.130** |
| E-Cad | Myo | 2872.24 | 0.638 | 1.261 | **393.77** | **0.557** | **0.882** |
| Jub | Myo | 1861.57 | **0.920** | 1.700 | **415.48** | 0.976 | **1.379** |
| Jub | Myo | 4253.36 | 1.389 | 2.129 | **1156.77** | **1.207** | **1.714** |

Since 4DR-GAN has a similar network depth and layer arrangement with Pix2Pix, the results show that predicting protein localizations by incorporating 4D information helps improve the quality of prediction in the view of 2D. The FID with I3D(z) and I3D(t) further evaluates the volumetric consistency and temporal consistency, respectively, and in most cases, 4DR-GAN outperforms Pix2pix. As expected, 4DR-GAN records a substantial improvement in temporal consistency, because the temporal correlation is ignored in Pix2Pix. For example, in the case of predicting E-Cad from Myosin, the score of volumetric consistency improves by 10.63%, from 0.762 to 0.681, and the score of temporal consistency improves even more, by 30.03%, from 1.612 to 1.130.

The experiment result demonstrates that when two functionally related proteins are correlated in 4D, 4DR-GAN incorporates the information in all four dimensions and achieves high-quality prediction.

### 3.2 PLP accurately recapitulates protein localization at subcellular levels

Because the function and activation state of the protein determine the subcellular localization, we further evaluated the similarity between the prediction and target ground truth, using key biological characteristics of the subcellular localization. Our data sets recorded the ventral cells of fly embryos (Fig. 2a, light purple) during a period when these cells turned from a flat sheet into a tube-like structure. This tissue shape change is driven by the combined action of Myo and E-Cad (Fig. 2a). Myo is a molecular motor that generates the contractile physical force that changes cell shape while E-cad connects Myo filaments in individual cell into tissue-level network^32,33^. During the imaging period, increasing the amount of Myo proteins are activated in ventral cells and the activation is restricted on the apical surface of the cell (Fig. 2a, red filaments). In confocal microscopy images, the activated Myo complexes are visualized as filamentous networks of high concentration, which appears in top slides of the image stacks (Fig. 2a and 2c, top row input). The inactive pool of Myo appears to be uniform and at low concentration since they diffuse freely inside the cell. To apply the force generated by active Myo to the cell, Myo network is connected to the cell membrane through the interaction with E-Cad complexes. E-Cad complexes provide adhesions between neighboring cells. In the images inactive E-Cad proteins uniformly diffuse on and label cell membrane with low intensity, whereas the E-Cad proteins engaged in cell adhesion are assembled into higher-order complexes and appear to be high-intensity clusters along the cell-cell boundaries (Fig. 2a; 2c, second row, target, 8×8 cells). By connecting to these E-Cad clusters, Myo filaments pull cell boundaries towards the center of the apical surface, therefore reducing cell apical surface areas (Fig. 2a). Meanwhile, in response to the force experienced by the E-Cad complex, Jub is recruited to the cell adhesion complex and detected as spots overlapping with a portion of E-Cad clusters along cell-cell boundaries^34^.

**Fig. 2.**
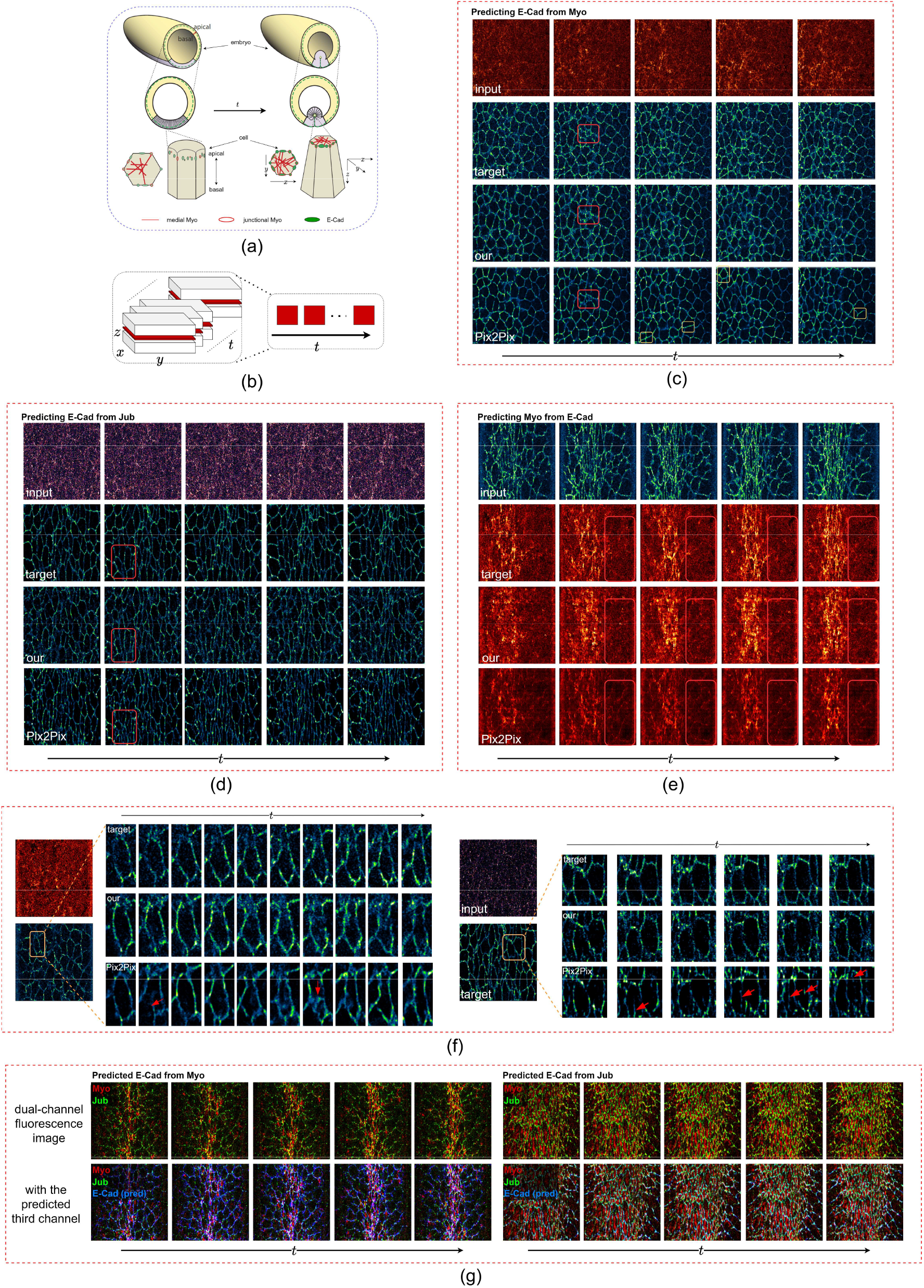
The subcellular localizations of the predicted proteins. (a) The known relationship between Myo and E-Cad. (b) The way of demonstrating 4D images with a z-slice and multiple time frames. (c) Localization prediction from Myo to E-Cad shown with the input Myo, the ground truth Ecad localization, our 4DR-GAN prediction results, and the Pix2Pix prediction results. The red boxes show 4DR-GAN predicts better cell outlines and the yellow boxes show the prediction results are too smooth without showing E-Cad clusters. (d) Localization prediction from Jub to E-Cad. The red boxes show Pix2Pix predicts inaccurately resulting in extra cell boundaries. (e) Localization prediction from E-Cad to Myo. The red boxes show that Pix2Pix predicts inaccurately resulting in visible cell boundaries. (f) Temporal consistency of prediction results. Red arrows show inconsistent predictions in time. (g) Generating an additional channel that cannot be imaged together. Best view with zoom in.

To evaluate the subcellular localizations of the predicted proteins, we picked a *z* slice close to the apical surface including major Myo and E-Cad signals and demonstrated the changes in consecutive *t* frames (Fig. 2b).

First, we compared the morphology of the protein localizations. Fig. 2c-e show the PLP results, with four rows displaying the input protein localization, the ground truth (GT) of target protein localization, and the prediction by Pix2Pix and our 4DR-GAN, respectively. As discussed above, E-Cad signals largely label cell boundaries. Consistent with the GT, 4DR-GAN produces correct outlines of individual cells, whereas extra or missing cells are often present in the Pix2Pix prediction (Fig. 2c, red rectangle). 4DR-GAN is also better at recapitulating the clustering behavior of E-Cad proteins. Because inactive E-Cad uniformly labels cell membrane and active E-Cad form clusters, the cell outlines visualized by E-Cad are dotted lines like the target ground truth. By comparison, the Pix2Pix results tend to be smooth lines without clusters (Fig. 2c, yellow rectangle). This shows that 4DR-GAN predicts the localization of activated E-Cad better than Pix2Pix highly possibly because 4DR-GAN utilizes the temporal information and allows capturing more information of the predicted proteins. This is especially important when input images are of low signal-to-noise ratios, such as Jub channel co-imaged with E-Cad.

As shown in Fig. 2d, the results produced by Pix2Pix show an extensive amount of extra cell boundaries that do not exist in the ground truth (Fig. 2d, red rectangle). In contrast, our 4DR-GAN prediction is able to generate correct cell boundaries for most cells. 4DR-GAN also generates more faithful predictions of Myo from E-Cad channel. 4DR-GAN is able to recapitulate both the active Myo pool (high-intensity network) and the inactive pool (uniform at low intensity). Among the active Myo, the majority localize in the center of the apical surface (medial Myo), while a minor pool localizes to some E-Cad complexes (junctional Myo). In Pix2Pix results, the predicted Myo shows lower intensities overall than GT and 4DR-GAN prediction. Interestingly, this inaccurate prediction affects medial Myo and inactive Myo more than junctional Myo, which results in a lower intensity ratio between medial Myo and junctional Myo in the Pix2Pix prediction. In addition, the localization of junctional Myo excessively resembles that of the input E-Cad rather than the ground truth Myo: cell outlines are clearly visible in Pix2Pix predicted Myo images even though cell outlines are barely visible in Myo images from GT and 4DR-GAN prediction (Fig. 2e, red box). These analyses show that 4DR-GAN gives rise to more accurate prediction of protein subcellular localization.

Secondly, we evaluated the temporal consistency of the predicted signals. Our 4DR-GAN is temporally more stable in terms of both pixel intensity and object morphology (Fig. 2f). In the ground truth and 4DR-GAN images, the pixel intensities of cell boundaries labeled by E-Cad are consistent between time frames. Whereas, in the predictions of Pix2Pix, the intensity of cell boundaries changes drastically, with some cell boundaries jumping from low to high intensity in a single time interval, only to drop in the next. Morphologically, it is observed that the shape of the same cell often changes sharply and cell boundaries can suddenly appear or disappear between time frames (red arrows in Fig. 2f). These predictions are clearly wrong as the shape and existence of cell boundaries do not change this drastically with our 10-sec frame rate. 4DR-GAN reduces these problems and maintains the temporal consistency of cell morphology.

Lastly, to test the effectiveness of generating an additional channel that cannot be imaged together, we applied the trained 4DR-GAN to dual-channel data sets of Myo and Jub and generated E-Cad images as the third channel. Either Myo channel or Jub channel can successfully predict E-Cad channel, as shown in Fig. 2g. The predicted E-Cad gives rise to cell boundaries that not only are of appropriate sizes and shapes but also cover Jub signals, consistent with Jub localizing exclusively to a portion of the E-Cad complex. Lastly, consistent with the known spatial relationship of Myo and E-Cad localization, Myo appears mostly inside the cell boundaries labeled by predicted E-Cad.

### 3.3 PLP predicts protein localization dynamics with high fidelity

Protein subcellular localization is dynamic during development and can change dramatically. The quality of dynamics of 4DR-GAN predicted protein localization is evaluated using the Myo channel predicted from the E-Cad channel since Myo subcellular localization changes in all five dimensions during the live imaging periods.

First, we compared Myo intensity between GT and prediction in Z and T dimensions (Fig. 3a). Similar to GT, in 4DR-GAN predicted channel, high-intensity Myo is only detected on the apical surface (the first several z slices) and becomes more and more intense during the imaging time frame. This is true for both medial and junctional Myo. Although the Pix2Pix prediction follows a similar pattern, the intensity of predicted Myosin is lower. This is especially prominent for medial Myo, consistent with the 2D analysis that finds a lower medial: junctional Myo ratio (Fig. 2e). This becomes clearer when analyzing the intensity profiles along Z at a given time point or the intensity increase with time at a given z (Fig. 3b and c). While GT and 4DR-GAN closely resemble each other, the profiles generated by P2P show lower intensities and somewhat deviated curves.

**Fig. 3.**
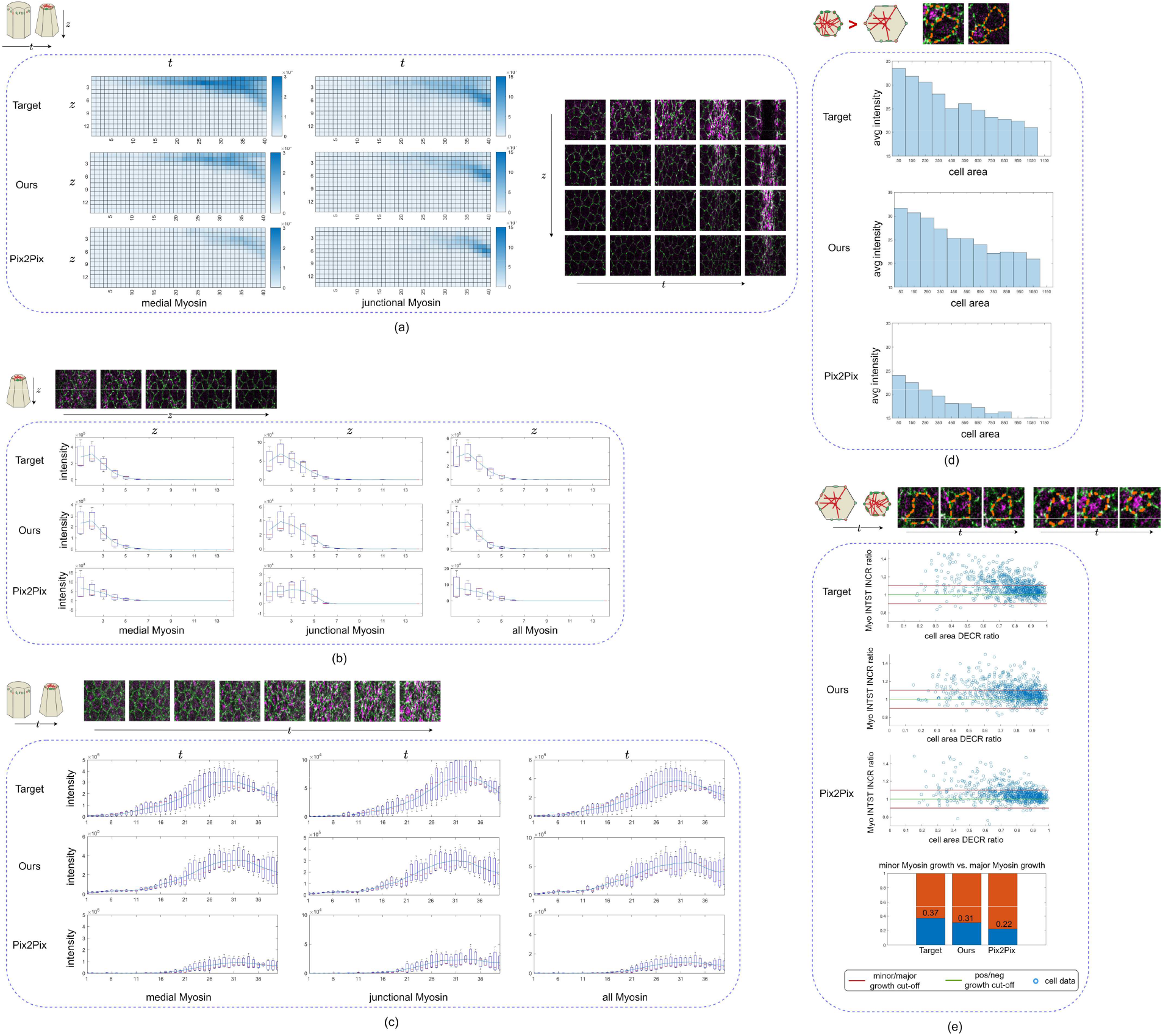
The localization dynamic of the predicted Myo from E-Cad. (a) The predicted Myo intensity in Z and T dimensions compared with GT. Medial and junctional Myo signals are separately reviewed because they have differential relationships with the input E-Cad. (b) The predicted Myo intensity dynamics in Z dimension with a fixed time frame (t=30) compared with GT. The example images show the GT signal in Z dimension. (c) The predicted Myo intensity dynamics in T dimension with a fixed z-slice (z=4) compared with GT. The example images show the GT signal in T dimension. (d) The relationship between cell area and average Myo intensity in prediction results compared with GT. The example images demonstrate that the cells with smaller apical surface areas are more likely to have higher active Myo. (e) The relationship between cell area decreasing (DECR) ratio and average Myo intensity (INTST) increasing (INCR) ratio. The example images demonstrate that when the cell apical surface area decreases continuously, the cell is more likely to have increasing active Myo. Best view with zoom in.

Secondly, we examined whether the predicted Myo localization is consistent with its biological function. Higher concentration (intensity) of filamentous Myo is correlated with higher contractile force^35^. By connecting to E-Cad complex, the contractile force reduces apical surface area. Therefore, at a given time point, cells with smaller apical surface area are more likely to have more active Myo. Fig. 3d quantifies the average Myo intensities for cells of different sizes during the 10 consecutive time points when Myo is extensively activated. These histograms show that Myo is indeed at higher levels in cells of smaller size. The histogram profile of 4DR-GAN prediction is closer to that of GT than the Pix2Pix result. Similarly, a given cell usually has more active Myo when its area is reduced. Fig. 3e shows the changing rate of Myo in cells of decreasing sizes at three time points. Within this short time frame (30 sec), 4DR-GAN prediction and GT have over 30% of cells that show more than 10% increase in Myo. This parameter in Pix2Pix prediction is 22% which is 30-40% less than that of 4DR-GAN prediction and GT.

### 3.4 Digital activation and inactivation predict correct consequences of protein loss-of-function and gain-of-function

The above analysis shows that, by integrating the information from all four dimensions, 4DR-GAN can accurately predict protein subcellular localizations and concentrations. Protein localization and concentration are not only the input and output of PLP, but are also closely related to protein activation states. For example, a higher concentration of Myo indicates more activated Myo proteins and correlates with higher physical tension generated by Myo. Therefore, we reason that it is possible to digitally control protein activities, through altering their localization and concentration in the images. This inspired us to develop effective digital activation and digital inactivation methods to digitally manipulate protein activities and predict functional consequences. The predicted channel of 4DR-GAN-based PLP should respond to the input change in a way consistent with their functional relationship.

To test the effectiveness of these digital operations, we first used Myo as the input channel and the change in cell apical surface area caused by Myo activities. We performed DI of Myo in the circled region by erasing Myo signals (Fig. 4a, second row). Since Myo produces contractile tension to reduce the cell apical surface, removing active Myo should lead to the relaxation of apical surface area, which appears in the image as bigger cells outlined by E-Cad. Indeed, an immediate response of the predicted E-Cad around the Myo knockout region is labeled: the cell outline marked by E-Cad in new prediction becomes bigger (red) than those in the prediction before the digital inactivation (cyan). This is consistent with observations from biological loss-of-function experiments where breaking Myo filaments with high power lasers leads to relaxation and expansion of the cell apical surface^33^. Fig. 4b shows the effect of DA of Myo by increasing Myo intensity in the circle, which represents an increase in the contractile force and should lead to a reduction in the cell apical surface area. The cell outline marked by E-Cad in the new prediction becomes smaller (red) than that in the prediction before the digital activation (cyan). Again, this is in line with the observations from biological experiments^36^, where forced Myo activation using genetic approaches induces cell apical surface area reduction.

**Fig. 4.**
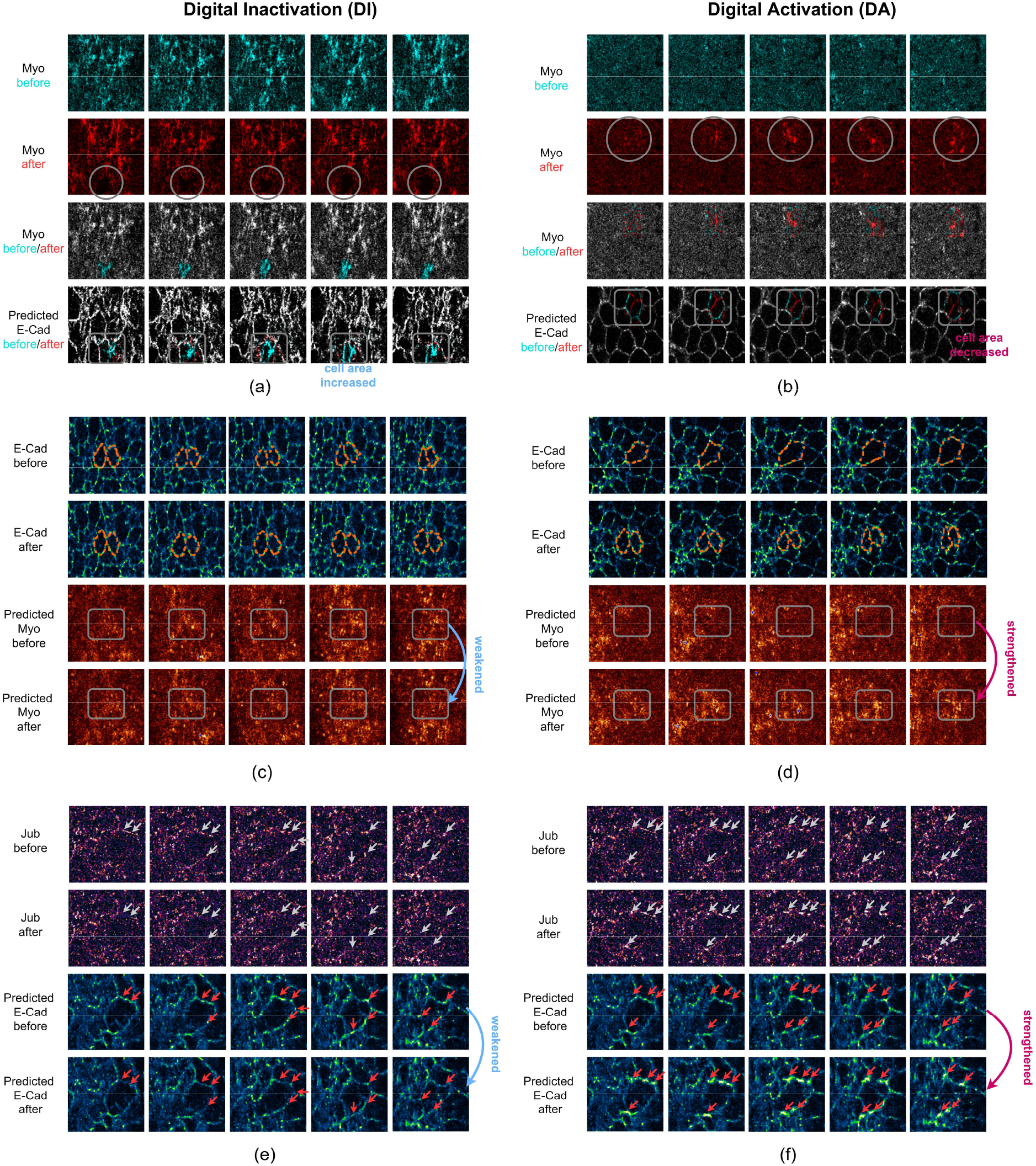
Digital inactivation and digital activation results. (a) Observing E-Cad and cell response when digitally inactivating Myo subcellular localization. (b) Observing E-Cad and cell response when digitally activating Myo subcellular localization. (c) Observing Myo response when digitally inactivating E-Cad and stopping apical areas shrinking. (d) Observing Myo response when digitally activating E-Cad and increasing apical areas shrinking. (e) Observing E-Cad response when digitally inactivating Jub. (f) Observing E-Cad response when digitally activating Jub. The gray circles and arrows highlight the DI or DA manipulation, the orange dotted lines show the apical surface areas manipulation, and the gray rectangles and red arrows highlight the responses. Best view with zoom in.

In the above case, Myo is the driving force to cause cell apical area to change. Next to ask, when we digitally manipulate the consequence, whether we can predict the localization pattern of the required Myo. Specifically, we tested whether keeping cells from decreasing their apical areas would decrease the predicted Myo intensity and whether forcing cells to shrink would increase the predicted Myo intensity. Both operations generated the expected results around the digitally altered regions. In the result, compared with the prediction without manipulation, Myo is weaker when cell areas are kept from being reduced (Fig. 4c), while Myo increases when one large cell is digitally split into two smaller ones that shrink in apical area (Fig. 4d).

To test whether this approach can be applied to a variety of proteins, we digitally activated and inactivated Jub and observed how E-Cad responds (Fig. 4e-f). Jub diffuses in cytoplasm when inactive. It can be activated in response to Myo-generated tension and recruited to E-Cad clusters^34^. This recruitment of Jub into clusters is hypothesized to stabilize cell-adhesion provided by E-Cad complexes^37^. Based on Jub localization properties, we digitally activated or inactivated Jub by strengthening or weakening Jub cluster intensity in the input images. It is observed that a weakened Jub cluster is translated into a weakened E-Cad cluster, and the two clusters overlap with each other. On the other hand, a Jub cluster of increased intensity is translated into a higher-intensity E-Cad cluster (Fig. 4e-f). This indicates that Jub and E-Cad not only colocalize in the images due to their molecular interaction, but there is also a strong correlation between Jub and E-Cad clusters’ intensity, consistent with Jub’s role in stabilizing E-Cad-based cell adhesion^34,37^. This also shows that DA and DI are versatile approaches applicable to proteins with a network-like localization (Myo) and proteins with a cluster-like localization (E-Cad and Jub).

## 4. Discussion

Compared with Pix2Pix, protein localization prediction generated by 4DR-GAN is more accurate in subcellular localization, temporal consistency, and dynamics. The experiment results demonstrate the importance of incorporating information from all spatial and temporal dimensions in the prediction of protein localization, which allows 4DR-GAN to capture more relationship features between two protein localizations. Noticeably, there are often different pools of the same protein that change with time and show differential localizations. When the pools of a protein play different roles in protein relationships, taking advantage of four dimensions simultaneously is the key for accurate prediction. For example, predicting E-Cad from Myo localization requires the network to differentiate between junctional Myo and medial Myo. 4DR-GAN is able to do so and accurately predict the localization of E-Cad when junctional and medial Myo signals change dramatically with time. Similarly, the prediction from Jub to E-Cad requires 4DR-GAN to learn the relationship between Jub, activate E-Cad (high-intensity clusters), and inactivate E-Cad (low intensity uniform membrane). These experimental cases suggest that 4DR-GAN can learn complex spatial and temporal relationships.

The proposed PLP method will benefit a variety of fluorescence microscopies, especially live imaging where fluorophore choices are limited. For example, in our experiments, imaging Jub and E-Cad together is already challenging due to low in vivo protein concentration and fluorophore limitations, let alone imaging three proteins. With PLP, we successfully predicted E-Cad from the imaged Myo channel in Myo-Jub data sets, which leads to high-quality signals for all three proteins. PLP can also be instrumental in case of hardware limitations such as the availability of laser lines and detectors on a microscope by predicting other channels from imaged channels.

Based on 4DR-GAN-based PLP, DA and DI are two novel tools we propose for protein functional relationship study. A key feature of DA and DI is the capacity to precisely manipulate a protein localization in space and time. When doing so, DA and DI reflect the immediate effect on the output protein localization and therefore shed light on the protein relationships locally and globally. The experimental results not only demonstrate that DA and DI can predict the correct consequences of protein loss-of-function and gain-of-function, but also suggest that the 4DR-GAN-based PLP learns correct protein relationships. DA and DI require manipulation designs appropriate for protein functions. For unknown protein functions, multiple designs should be considered and analyzed.

Another key feature of DA and DI is that the outputs respond to input manipulation regardless of causality between proteins or the structure labeled by the protein. When the upstream protein is manipulated digitally, it mimics biological loss-of-function and gain-of-function experiments and the downstream protein responds in the predicted localization. However, when manipulation is applied to the downstream protein, while the upstream protein does not respond in biological experiments, it does in digital experiments. Specifically, when the downstream protein is digitally manipulated into a certain localization, the prediction provides a clue on the localization of the upstream protein localization that is required to drive the downstream protein into this certain localization. Therefore, DA and DI provide additional information in such cases. Overall, the realization of DA and DI provides a convenient and low-cost way to study protein functions and relationships and guides experimental designs in biological studies of unknown proteins.

Besides DA and DI, there are other possible applications of PLP. For example, in the cases of predicting the localizations of E-Cad from that of Myo and backward, we observe that the prediction of E-Cad from Myo has better perceptual quality than the prediction of Myo. The high prediction quality of E-Cad could imply that Myo is a major factor affecting E-Cad localization, while the information from E-Cad alone is insufficient and factors other than E-Cad contribute significantly to Myo localization. This observation also suggests potential future works of PLP. By adopting multiple informational sources, the performance of PLP for proteins that have multiple factors can be improved. In addition, analyzing the prediction quality may provide clues to the causality between proteins or cellular structures labeled by proteins.

## Acknowledgements

We acknowledge the support for this research: UNLV TTGRA and NIH Pathway to Independence Award (K99/R00 HD088764).

