## Supplementary for "Digitally Predicting Protein Localization and Manipulating Protein Activity in Fluorescence Images Using Four-dimensional Reslicing GAN"

### Network implementation

**4DR-GAN.** The proposed 4DR-GAN consists of a generator *G* and a discriminator *D*. *G* is an encoder-decoder network that has two encoders and a decoder. The layer arrangement of the encoders and the decoder in 4DR-GAN is similar to that of Pix2Pix. Fig. 5 demonstrates the implementation details of *G* and *D*. Particularly, the stride of convolutional layers in *G* is asymmetric to preserve the feature size on z-axis and t-axis. Subsequently, the feature maps with an identical shape can be resliced and paired. The decoder of *G* has two feature map composing methods: concatenation and residual SPADE (Res SPADE)[38]. The concatenation layer is used in most cases, whereas we find Res SPADE benefits when the boundaries in the input are important to the prediction (e.g. Ajuba and E-Cad), because Res SPADE was proposed for semantic boundary refinement.

In the section of results, we introduced the structure of 4DR-GAN with feature reslicing for the mainstream. Here, Extended Data Fig. 1 demonstrates the feature reslicing details for mainstream and skip connections.

**Pix2Pix.** Pix2Pix[25] is the baseline network we compared with. The original Pix2Pix is implemented in 2D, whereas we expanded it to 3D by replacing all 2D network layers with 3D layers. In this way, Pix2Pix is able to incorporate localization information in three dimensions and therefore is optimized for PLP for 4D images.

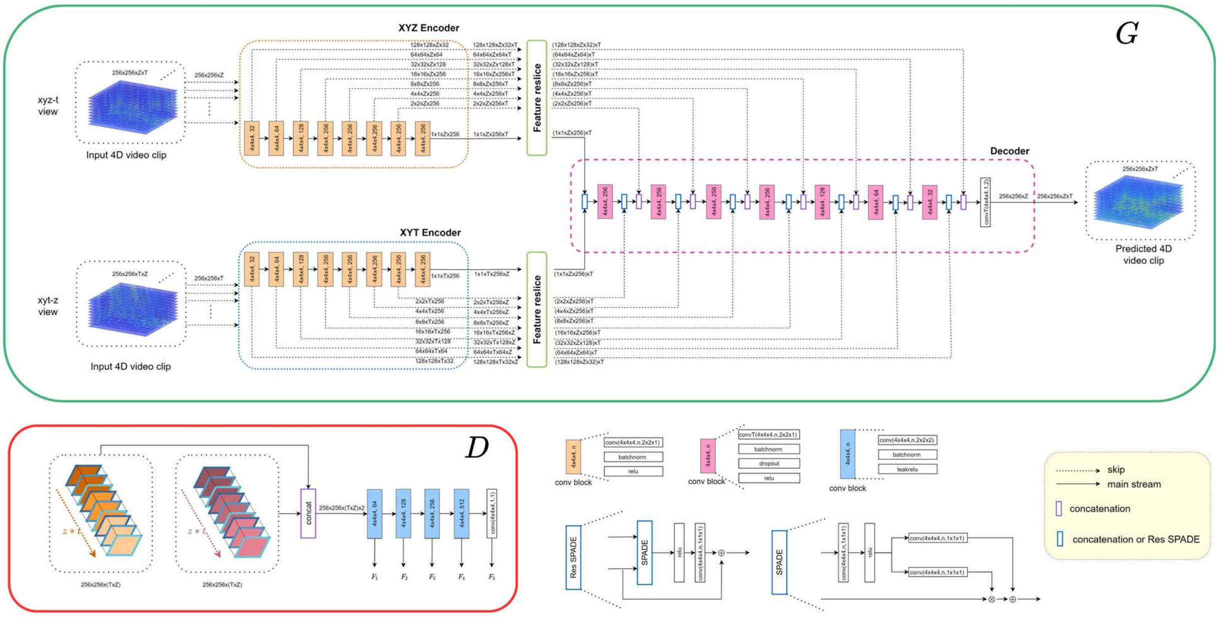


Extended Data Fig. 1 | 4DR-GAN architecture. In our experiment, the input of G is 256x256xZxT, where Z and T are the sample sizes in the z-axis and t-axes, respectively. The output of *G* has the same size. In particular, conv(4x4x4,n,2x2x1) represents a 3D convolutional layer with a 4x4x4 filter size, n filters, and an asymmetric stride of 2x2x1. Similar to Pix2Pix, the first conv block in encoders works without batch normalization layers. As noted, we use ReLu activation in our generator implementation rather than LeakReLu used in Pix2Pix. D is a PatchGAN[25] that takes two channels as inputs in our implementation. The two inputs to *D* are concatenated to extract features on protein localization, dynamics, and interaction. The feature maps from each convolutional block are exported for loss calculation. All adding and multiplication operations are element-wise.

### Training objective function and parameters

To tune the generator and the discriminator together, we combined the adversarial loss, feature loss, and reconstruction loss in the training objective to achieve stable training and realistic prediction. If we denote the input 4D image as $V_\alpha$, the target as $V_\beta$ with the data distribution $\mathbb{E}_{(V_\alpha,V_\beta)}$, the prediction is $\widehat{V_\beta} = G(V_\alpha)$. To tune the generator *G* and discriminator *D*, we have the adversarial loss as

$$\min_G \max_D \mathcal{L}_{GAN}(G,D) = \mathbb{E}_{(V_\alpha,V_\beta)}[\log(D(V_\alpha,V_\beta)) + \log(1 - D(V_\alpha,\widehat{V_\beta}))]$$

The feature matching loss based on the discriminator is adopted to stabilize the training, thus improving the GAN loss[39]. If $D^j$ describes the $j^{th}$ convolutional block in *D*, the output of $D^j$ is $F^j_{V_\alpha,V_\beta} = D^j(V_\alpha,V_\beta)$. Subsequently, the feature matching loss $\mathcal{L}_{FM}(G,D)$ is

$$\mathcal{L}_{FM}(G,D) = \mathbb{E}_{(V_\alpha,V_\beta)} \sum_{j=1}^{J} \frac{1}{N_j}[\left\| F^j_{V_\alpha,V_\beta} - F^j_{V_\alpha,\widehat{V_\beta}} \right\|_1],$$

where $J$ is the total number of *D*'s output used in loss calculation, and $N_j$ is the total number of elements in $F^j_{V_\alpha,V_\beta}$. $J = 1$ in our implementation.

Furthermore, we applied the reconstruction loss to minimize the pixel level difference between the predicted result and the target. As pointed out in existing works[25], L1 loss has advantages, including less blurring and high-quality distribution approximation:

$$\mathcal{L}_{L1}(G) = \mathbb{E}_{(V_\alpha,V_\beta)}[\left\| V_\beta - \widehat{V_\beta} \right\|_1]$$

The final combined objective function is:

$$\min_G[(\max_D \mathcal{L}_{GAN}(G,D)) + \lambda_1 \mathcal{L}_{FM}(G,D) + \lambda_2 \mathcal{L}_{L1}(G)],$$

where 1 and 2 are 100 in our implementation.

Both baseline and 4DR-GAN were trained on the same objective function for fair comparison. In the training process, the baseline took 3D fluorescence image at each time frame as the input with batch size 2. The baseline was trained for 100 epochs. 4DR-GAN was trained for 40 epochs because it took 4D images as the input with batch size 1. The optimizer used was Adam[40] with a learning rate of $2\text{x}10^{-4}$ with a decay rate of 0.96 every 1000 optimization steps. The network and the training program were implemented by Python and Tensorflow 2.3 on a computer configured with an Intel i7-8700k, 48GB RAM, and a NVIDIA Titan RTX with 24GB GRAM.

### Evaluation metrics and their implementation

Originally proposed for evaluating image generation, FID[27] adopts the pretrained InceptionV3[29] network as the feature extractor and removes the last classification layer. Subsequently, when gaining an image and sending it to the InceptionV3, FID obtains the features and calculates the mean and covariance matrix. In this way, it obtains the features from both the prediction and the target ground truth. The mean and the covariance matrix for the prediction and the target ground truth are denoted as $\tilde{\mu}$ and $\tilde{\Sigma}$, and $\mu$ and $\Sigma$, respectively. The FID formula is

$$FID = \|\mu - \tilde{\mu}\|^2 + Tr(\Sigma + \tilde{\Sigma} - 2\sqrt{\Sigma\tilde{\Sigma}})$$

In the work of video translation[31], the authors proposed a video-based FID using pre-trained video recognition networks for video generation evaluation to measure both visual quality and temporal consistency. In our work on 4D images, the original FID was used to measure the quality slice-wise, and the video-based FID was used to measure the volumetric and temporal consistency separately. Like the work[31], the InceptionV3 pre-trained on ImageNet and the I3D[30] pre-trained on Kinetics400 were used. To measure the volumetric and temporal consistency, for a 4D image, each t frame was seen as a z-axis video and each z-frame was seen as a t-axis video.

To segment medial and junctional Myo (Fig. 3a-c), we differentiated E-Cad and Myo by thresholding, as shown in Extended Data Fig. 2. For E-Cad, adaptive thresholding[41] was used and for Myo, fixed thresholding (thre=0.25) was used. The E-Cad and Myosin masks were used to locate junctional and medial Myosin. The junctional Myo mask was calculated by multiplying the E-Cad and Myo masks while the medial Myo mask was the multiplication of the reversed E-Cad mask and Myo mask. Applying the junctional and the medial Myo masks on Myo images produced the junctional and medial Myosin images respectively. In Fig. 3d-e, we used adaptive thresholding on E-Cad with image closing to segment cells, and we further excluded connected cells by detecting the high-level E-Cad intensity inside segmented cells, as shown in Extended Data Fig. 3. Furthermore, for changing rate analysis in Fig. 3e, we tracked cells in three time points to observe the mean intensity variation of Myo, with the tracking results shown in Extended Data Fig. 4. We used MATLAB to implement this method.

### Digital activation (DA) and digital inactivation (DI) practice

Two methods were implemented to apply digital inactivation (DI) on different proteins. For Myo, we covered the Myo signals with masks. Masks were drawn in each z slice at each time frame to cover the target Myo object. This operation was performed by using ImageJ[42]. For cell outlines labeled by E-Cad and Ajuba, the target cell in the first time point was copied to all later time points, using MATLAB.

To apply digital activation (DA) and mimic in vivo protein behaviors, we copied the images of activated proteins from other regions to the targeted region. A single cell was used as a unit for this operation. Cells with activated proteins were copied and passed to the target region to mimic activation in this region. This process was conducted in MATLAB.

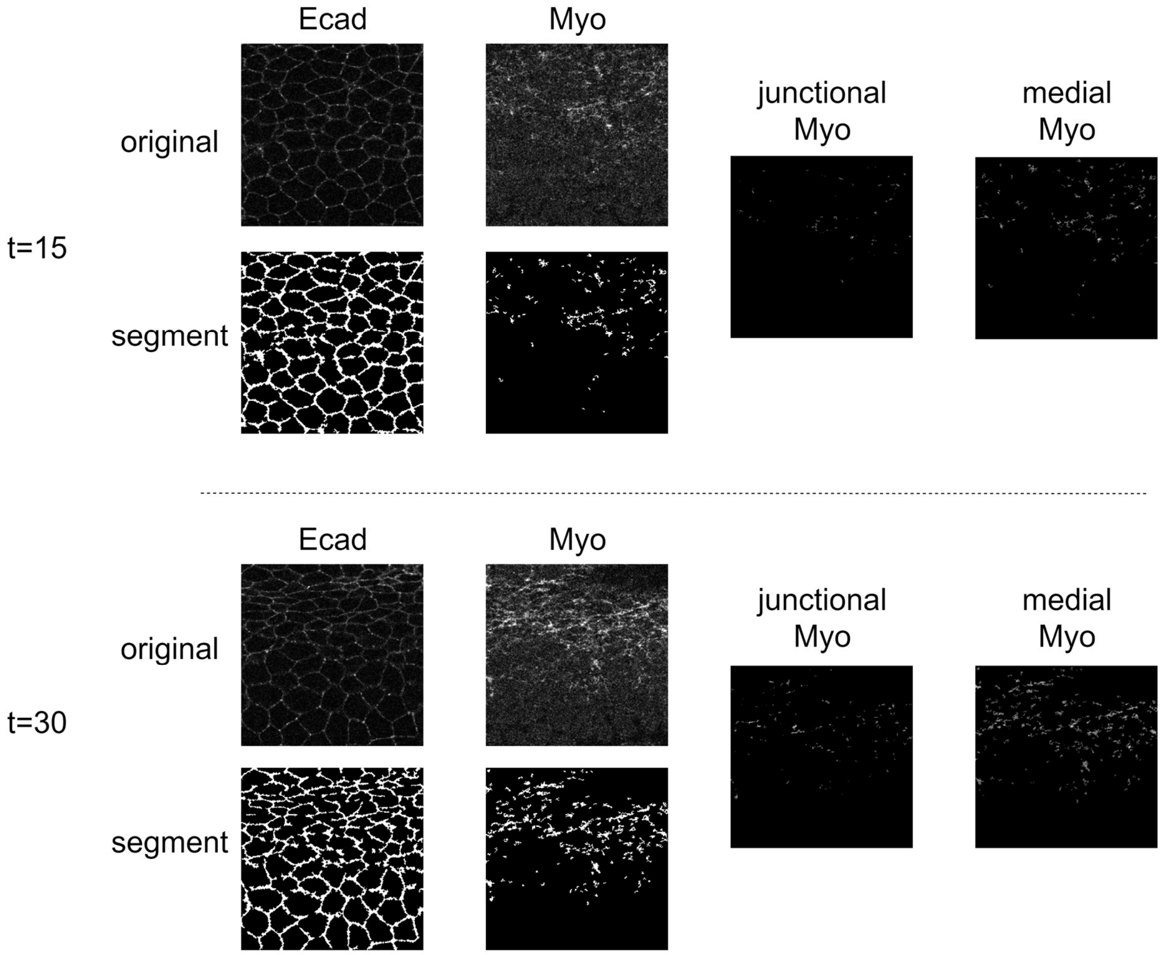


Extended Data Fig. 2 | Junctional and medial Myosin differentiation by E-Cad and Myo segmentation. For a 5D fluorescence image with total 40 time frames, we demonstrate two thresholding results of E-Cad and Myo on 15 and 30 time frames.

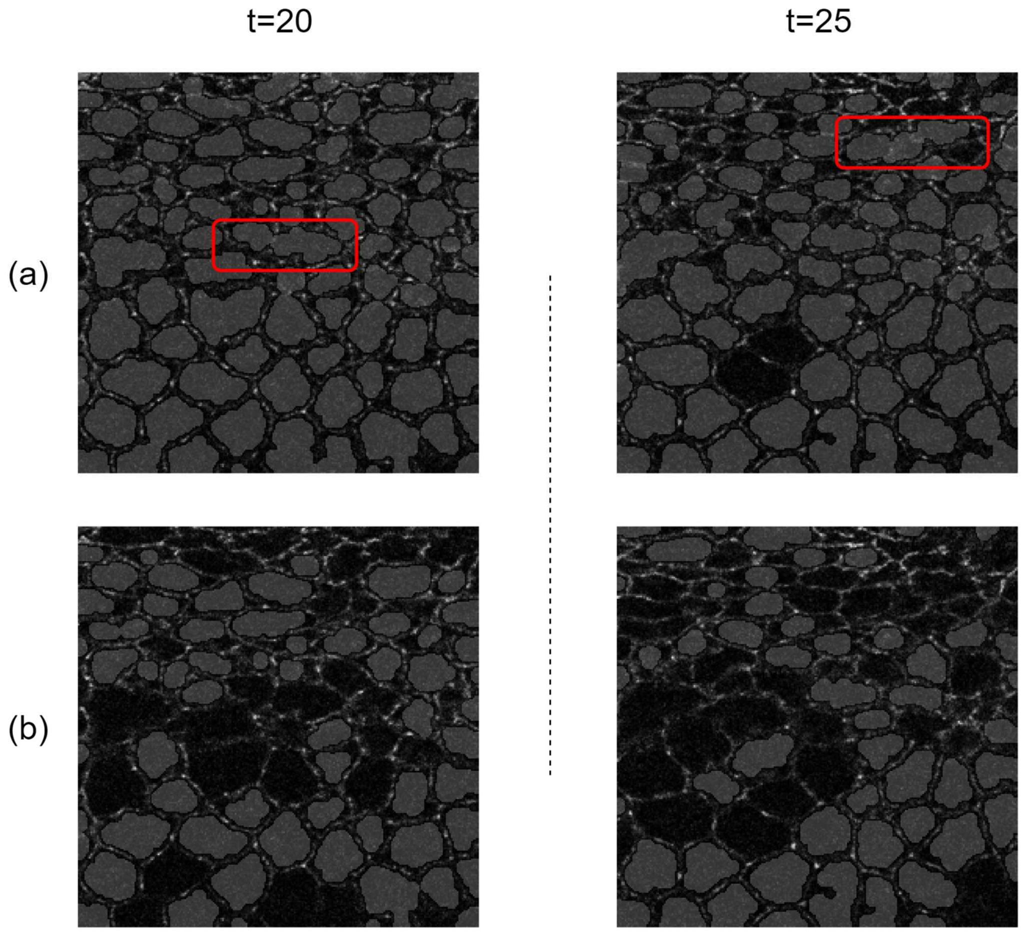


Extended Data Fig. 3 | Cell segmentation and quality control; (a) cell segmentation by reverse E-Cad segmentation; (b) segmentation quality control to exclude connected segmentations. The background is E-Cad localization which is used to define cell boundaries using thresholding. The gray masks are segmented cells. The quality control is excessively performed intentionally to fully exclude connected segmentations so that evaluations are performed precisely.

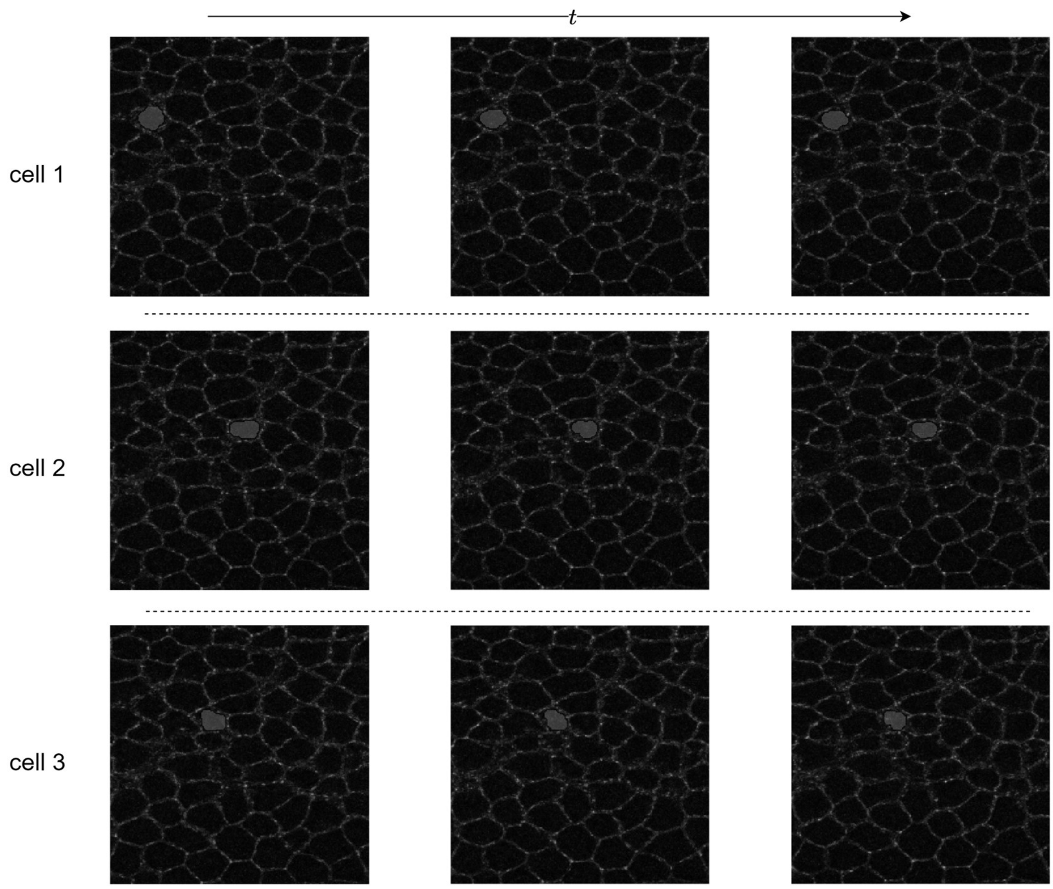


Extended Data Fig. 4 | Three examples of cell tracking based on cell segmentation. The gray masks in a row demonstrate successful cell tracking in three continuous time points. Cells that size decrease in three time points are used to analyze the relationship between size decrease and medial Myosin increase.

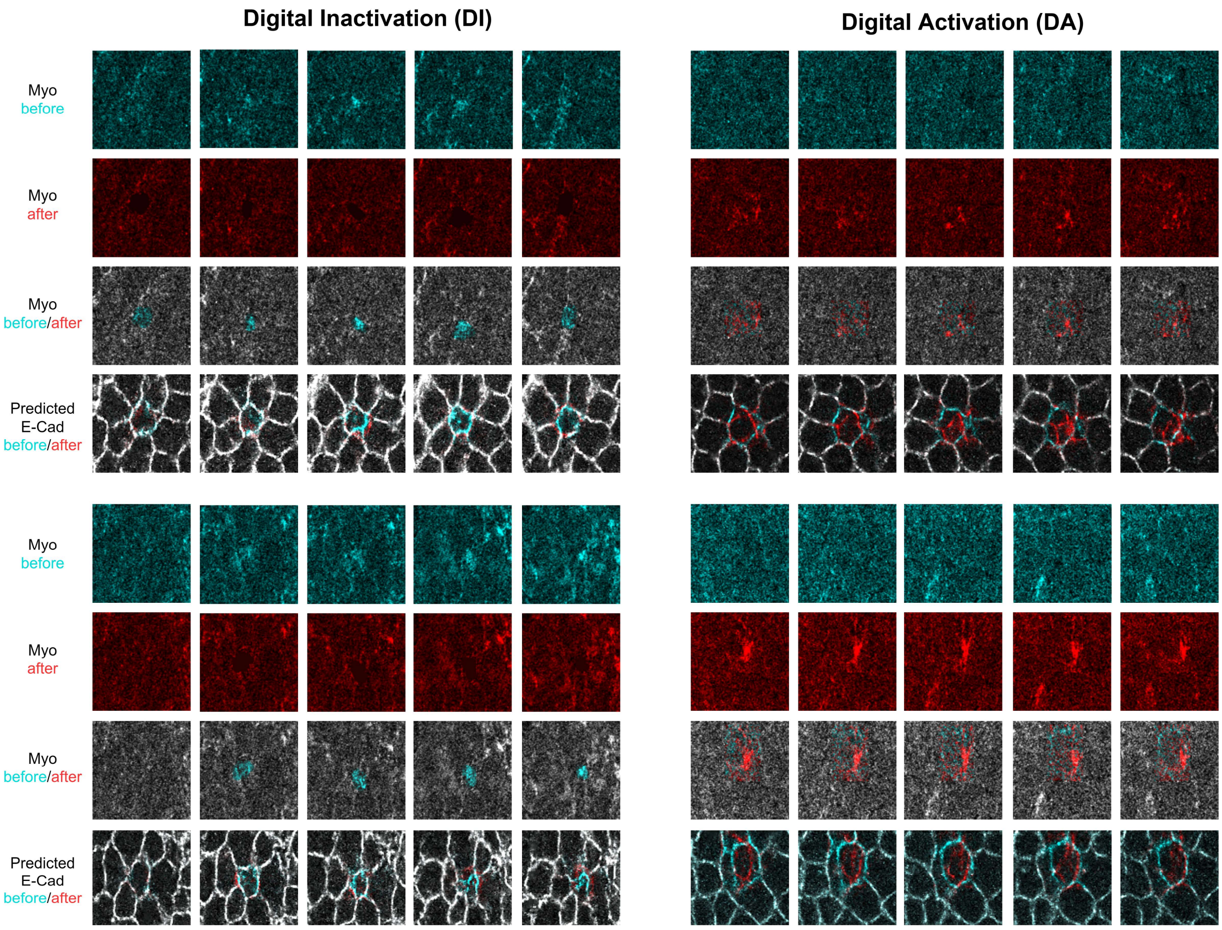


Extended Data Fig. 5 | Additional examples of DI and DA results on observing E-Cad response when manipulating Myo. The contrast of cyan and red shows the manipulation of Myo and the response of Ecad. Best view with zoom in.

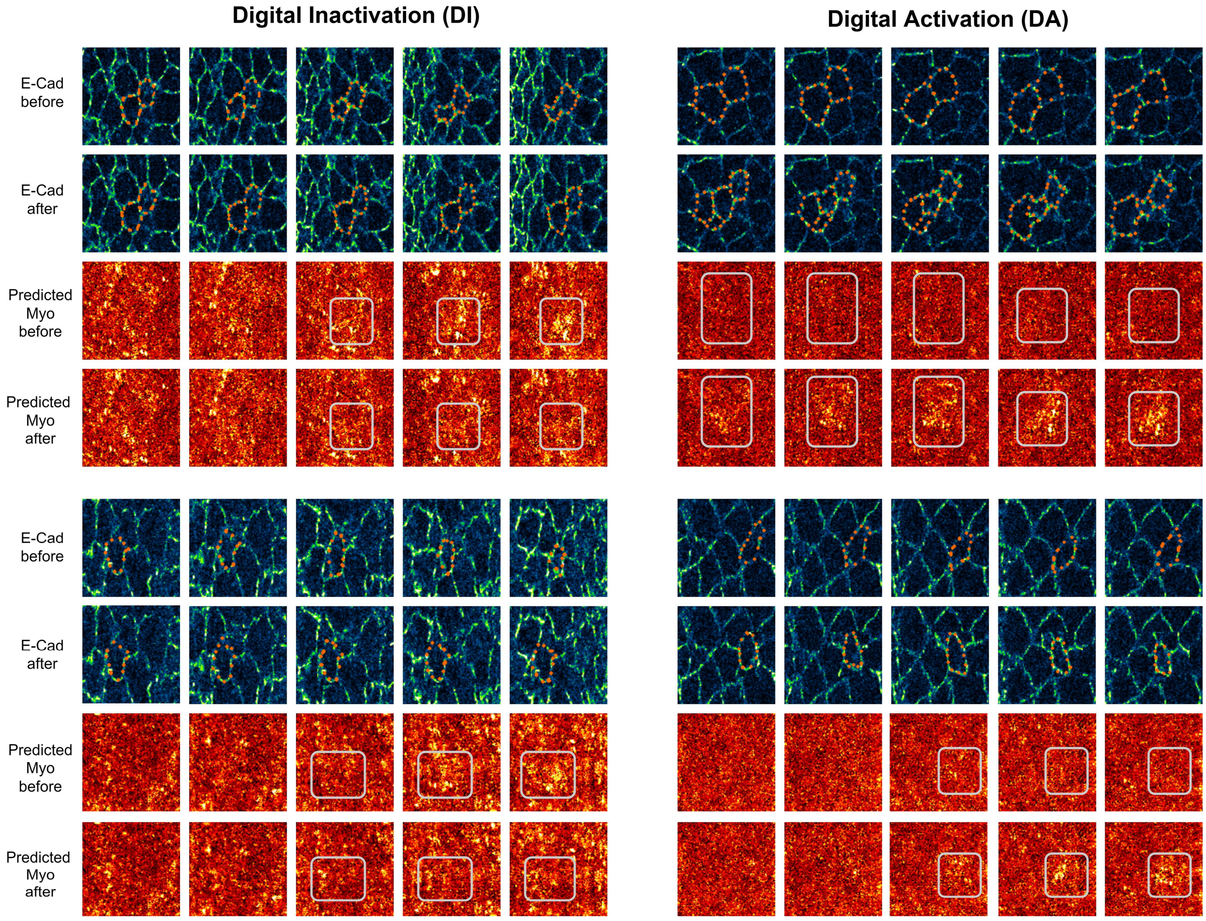

Extended Data Fig. 6 | Additional examples of DI and DA results on observing Myo response when manipulating E-Cad. The orange dotted lines highlight the apical surface area manipulation and the gray rectangles highlight the response. Best view with zoom in.

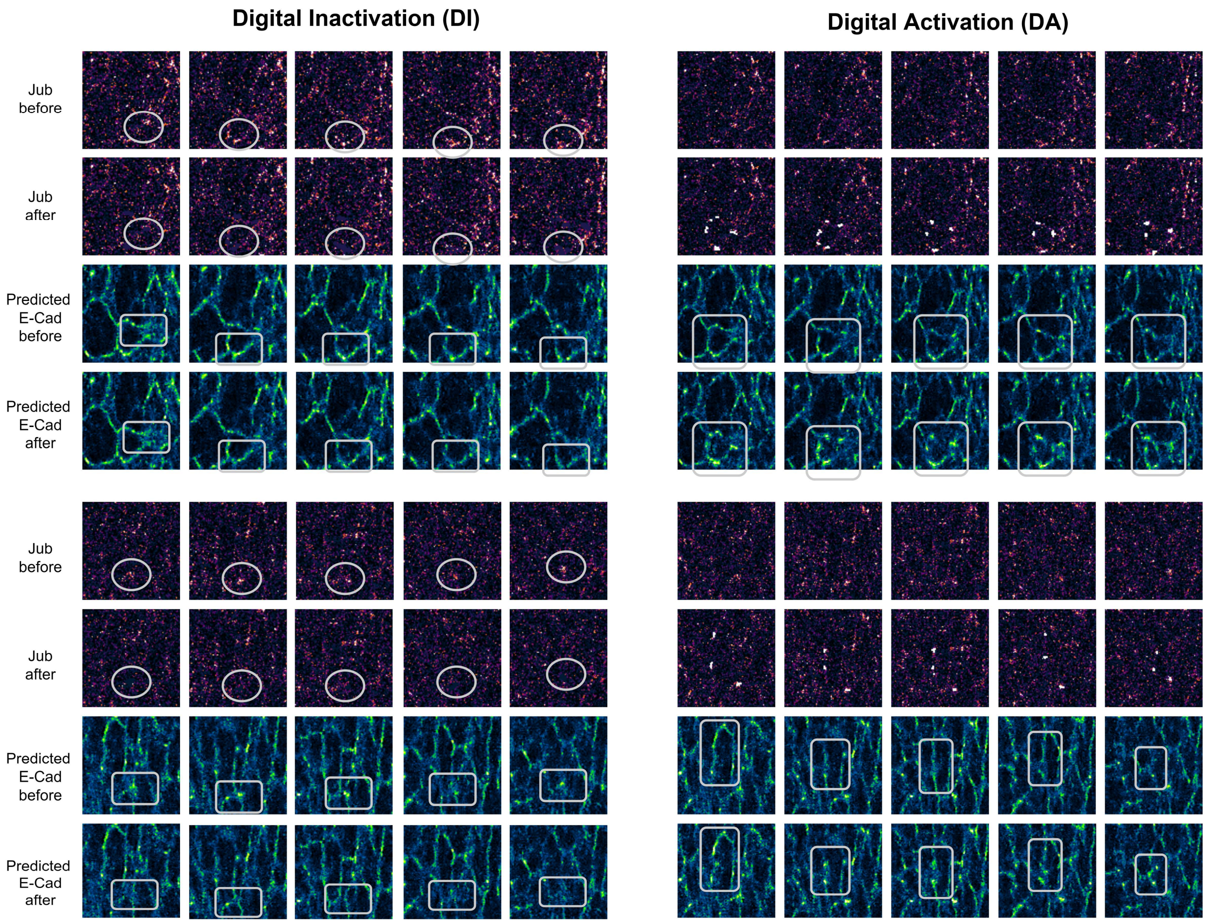


Extended Data Fig. 7 | Additional examples of DI and DA results on observing E-Cad response when manipulating Jub. The gray circles highlight the manipulation and the gray rectangles highlight the response. Best view with zoom in.
